# Topography, spike dynamics and nanomechanics of individual native SARS-CoV-2 virions

**DOI:** 10.1101/2020.09.17.302380

**Authors:** Bálint Kiss, Zoltán Kis, Bernadett Pályi, Miklós S.Z. Kellermayer

**Author notes:** These authors contributed equally (Bálint Kiss, Zoltán Kis).

## Abstract

SARS-CoV-2, the virus responsible for the current COVID-19 pandemic, displays a corona-shaped layer of spikes which play fundamental role in the infection process. Recent structural data suggest that the spikes possess orientational freedom and the ribonucleoproteins segregate into basketlike structures. How these structural features regulate the dynamic and mechanical behavior of the native virion, however, remain unknown. By imaging and mechanically manipulating individual, native SARS-CoV-2 virions with atomic force microscopy, here we show that their surface displays a dynamic brush owing to the flexibility and rapid motion of the spikes. The virions are highly compliant and able to recover from drastic mechanical perturbations. Their global structure is remarkably temperature resistant, but the virion surface becomes progressively denuded of spikes upon thermal exposure. Thus, both the infectivity and thermal sensitivity of SARS-CoV-2 rely on the dynamics and the mechanics of the virus.

**One sentence summary:** The native coronavirus 2 displays a dynamic surface layer of spikes, a large mechanical compliance and unique self-healing capacity.

## INTRODUCTION

Severe acute respiratory syndrome coronavirus 2 (SARS-CoV-2), the infective agent behind the current coronavirus disease (COVID-19) pandemic (Zhou et al., 2020; Zhu et al., 2020), is an enveloped ssRNA virus with a corona-shaped surface layer of spikes that are thought to play an important role in the infection mechanism (Hoffmann et al., 2020; Shang et al., 2020; Walls et al., 2020; Wang et al., 2020; Watanabe et al., 2020). Structural information about the spike protein has been acquired either on crystals of purified protein (Henderson et al., 2020; McCallum et al., 2020; Walls et al., 2020; Wrapp et al., 2020) or on fixed and frozen virus particles (Ke et al., 2020; Turonova et al., 2020; Yao et al., 2020). It has been suggested that the spike hinges provide structural flexibility (Ke et al., 2020; Turonova et al., 2020). High-resolution cryoelectron tomography observations indicate that the ribonucleoprotein (RNP) of SARS-CoV-2 is partitioned into spherical, basketlike structures (Yao et al., 2020). However, the surface dynamics and mechanical properties of native virions remain to be understood. Here we employed atomic force microscopy (AFM) and molecular force spectroscopy (de Pablo and Schaap, 2019; Kellermayer et al., 2018; Kiss et al., 2020) to investigate the topographical and nanomechanical properties of native SARS-CoV-2 virions immobilized on an anti-spikeprotein functionalized substrate surface. The unique single-particle approach revealed that the surface layer of spikes on SARS-CoV-2 is highly dynamic, the virion is unusually compliant and resilient, and it displays an unexpected global thermal stability.

## RESULTS AND DISCUSSION

### High-resolution topography of chemically fixed SARS-CoV-2

The topographical structure of individual SARS-CoV-2 virus particles bound to substrate surface was imaged by using AFM (**Fig.1.a**). To increase the efficiency and specificity of binding we employed a monoclonal anti-spike-protein antibody, which resulted in a nearly hundred-fold enhancement in the density of substrate-bound virions (**Fig. S1**). AFM images of glutaraldehyde-fixed SARS-CoV-2 revealed spherical virions (**Fig.1.b**) with somewhat variable dimensions (**Table S1**) and a rugged surface (**Fig.1.c**). The mean central height of the virions (**Fig. S2**), the structural parameter least sensitive to AFM tip convolution, was 62 nm (± 8 nm, S.D.) The height was smaller than the virion diameter measured in cryo-electron microscopic images (Ke et al., 2020; Turonova et al., 2020; Yao et al., 2020), suggesting that the virus particles were partially flattened on the substrate. The 3D-rendered AFM image (**Fig.1.d**) supported this interpretation and revealed that the rugged surface is due to the presence of protrusions which we identify as the spikes (S protein trimers). In high-resolution (pixel size 5 Å) AFM images (**Fig.1.e**) individual S trimers were resolved. Visual inspection of the S trimers pointed at their positional, rotational and flexural disorder in the viral envelope. The mean nearest-neighbor distance between the S trimers, and their topographical height were 21 nm (± 6 nm, S.D.) and 13 nm (± 5 nm, S.D.), respectively (**Fig. S3**, **Table S2**). From the mean nearest-neighbor distance and the virion dimensions we calculated that an average 61 spikes cover the SARS-CoV-2 virus particle surface. This number exceeds those reported recently (24 (Ke et al., 2020), 26 (Yao et al., 2020) and 40 (Turonova et al., 2020)), suggesting that the spike number is highly variable and may be regulated during virus assembly and maturation. The flexural disorder observed here supports the interpretation of cryo-electron microscopic data (Ke et al., 2020; Turonova et al., 2020; Yao et al., 2020), revealing a high degree of spike flexibility. We propose that the positional and rotational disorder of S trimers are due to their mobility in the virus envelope.

**Fig. 1.**
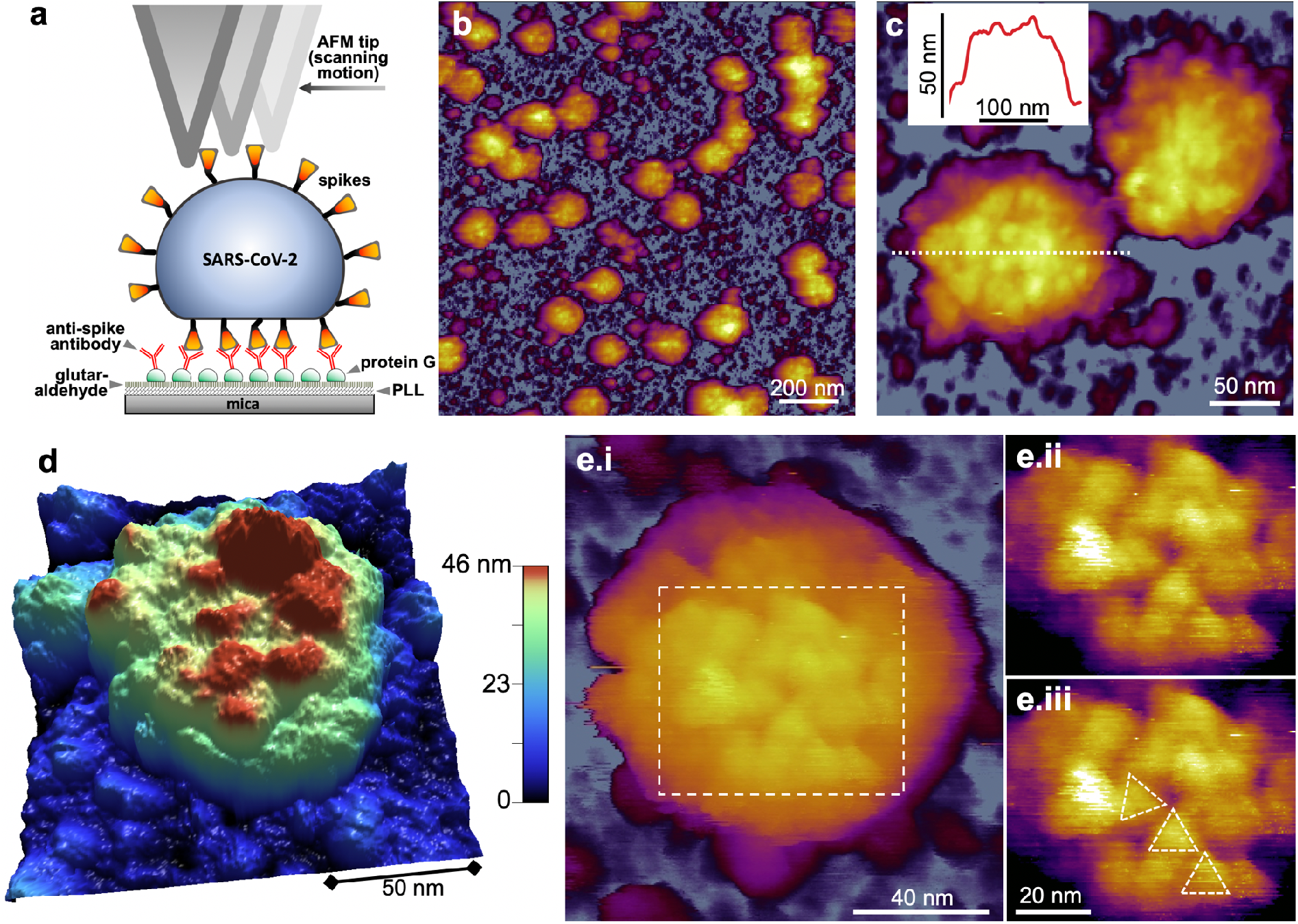
Topographical structure of SARS-CoV-2 virions treated with 5 % glutaraldehyde for structural preservation. **a.** Schematics of scanning substrate-surface-bound virions with the AFM tip. PLL: pol-L-lysine. **b.** AFM image of an overview (1.5 x 1.5 μm) sample area. **c.** Zoomed-in AFM image of SARS-CoV-2 virions. The virion surfaces are covered with protrusions that correspond to spikes (S protein trimers). **Inset**, topographical profile plot measured along the center of one of the virions (dotted line). The profile plot reveals a rugged surface. **d.** 3D-rendered image of a SARS-CoV-2 virion. A somewhat flattened virion is observed, pointing at a global flexibility of the virion. **e.** High-resolution AFM image of a SARS-CoV-2 virion displaying axial view of S trimers. **i.** AFM image of the entire virion. **ii.** Enlarged and contrast-enhanced image of the rectangular area. **iii.** The same AFM image with overlaid triangles indicating S trimer orientation. The spikes apparently display translational, rotational and flexural disorder owing to their flexibility.

### Structure and dynamics of native SARS-CoV-2

To circumvent the effects caused by chemical fixation and uncover the spike dynamics *in situ*, we investigated the topography of unfixed, native SARS-CoV-2 virions (**Fig.2**). Unexpectedly, we were unable to resolve S trimers on the virion surface at any of the investigated scanning strengths (**Fig. S4**); rather, the virus particles displayed a blurred, smooth surface (**Fig.2.a**). The mean central height of the native virions was 83 nm (± 7 nm, S.D.), which is significantly greater than that observed for the fixed (**Fig.2.b**). We interpret the blurring of virion topography as the result of aperture error caused by time averaging of spike movement within the sampling region of each image pixel, hence the increase in virion height is caused by the AFM tip scanning an apparent dynamic surface (**Fig.2.c**). Considering that the typical pixel sampling frequency (*f_s_*) in these images was 308 Hz, the frequency of spike dynamics exceeds the Nyquist frequency of *f_s_*/2 (154 Hz). Because the pixel size (*d*) is 2 nm, the spikes dynamically fluctuate, within a space limited by their flexible neck, with a speed (*v*=*df_s_*/2) exceeding ~0.3 nm/ms. Most plausibly, spike motion is dictated by the Brownian dynamics of the receptor-binding-domain (RBD) trimer which may then be thought of as a tethered particle. Spike mobility in the virus envelope may contribute further to the observed dynamics. An alternative explanation for the observed blurred virion surface is that the spikes evade the moving AFM cantilever tip which then scans the envelope surface. We exclude this possibility, however, because, while it relies on a similarly dynamic spike behavior, a reduced virion height should have been observed. We speculate that the rapid spike motion revealed by these experiments contributes to an efficient dynamic search by the virion on the surface of the targeted host cell, which explains why SARS-CoV-2 is at least as infective as the influenza virus (Petersen et al., 2020) in spite of its fewer spikes (up to ~60 in SARS-CoV-2 *versus* up to ~350 in influenza A (Harris et al., 2006)).

**Fig. 2.**
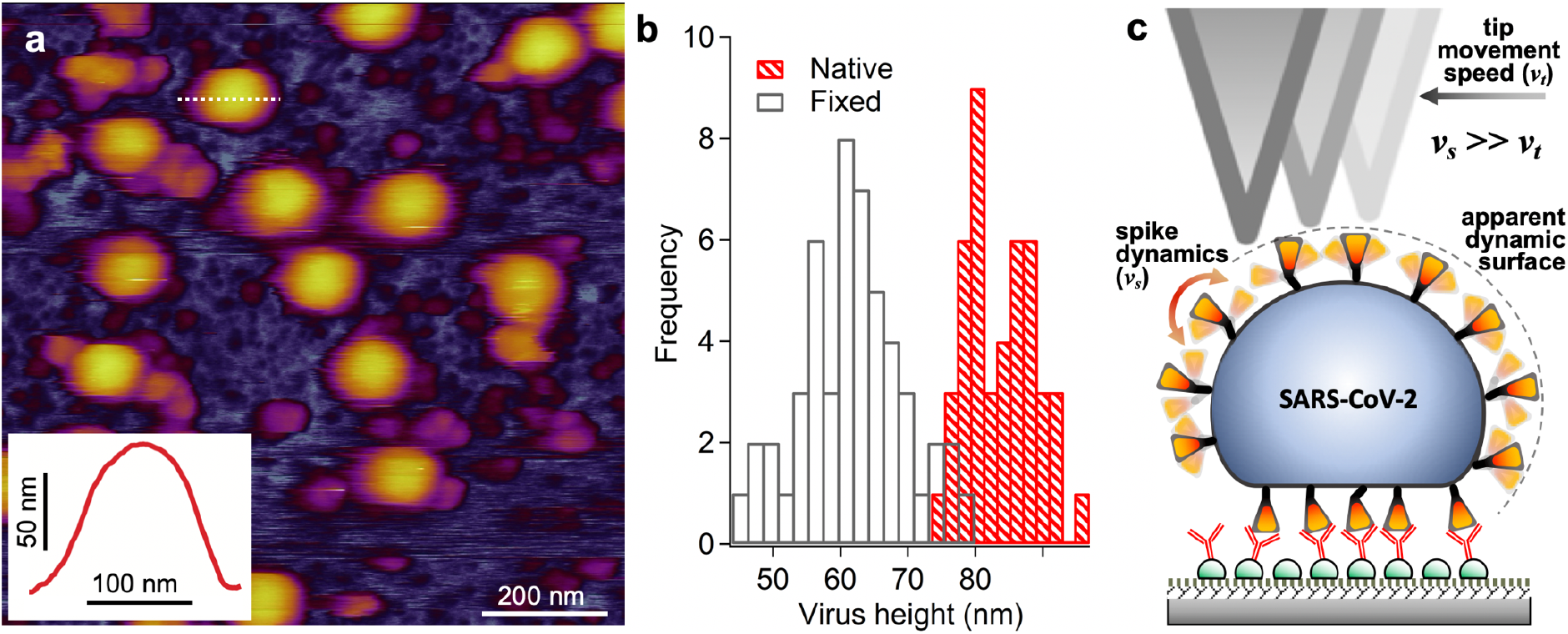
Topographical analysis of native, unfixed SARS-CoV-2 virions. **a.** AFM image of an overview (1 x 1 μm) sample area. Surface protrusions are not resolved, and a blurred, smooth topography is observed. **Inset**, topographical profile plot measured along the center of one of the virions (dotted line). The profile plot reveals a smooth surface. **b.** Distribution of the topographical maximal central height of fixed and unfixed SARS-CoV-2 virions obtained from particle analysis (see **Extended Data Fig.2**). Mean particle height (±S.D.) of fixed and unfixed virions are 62 ± 8 nm and 83 ± 7 nm, respectively. Unfixed virions have a significantly larger particle height than the fixed ones. **c.** Schematics explaining the dynamically enhanced height of the unfixed virion.

### Nanomechanics of SARS-CoV-2 virions

We investigated the mechanical properties of SARS-CoV-2 by lowering the cantilever tip on the vertex of individual virions selected on the AFM image (**Fig.3.a**). The virion was indented by pressing the tip downwards (**Fig.3.b**) with constant velocity (typically 0.5 μm/s) until a preset maximum force, measured by the cantilever deflection, was reached (typically 2-3 nN). Such a nanomechanical manipulation did not result in permanent topographical changes (**Fig.3.c**) in spite of completely compressing the virion so that the tip reached all the way to the substrate resulting in a wall-to-wall deformation (**Fig.3.d**). In the initial stage of indentation, immediately following the landing of the tip on the virion, we observed a linear force response that allowed us to measure virion stiffness (**Table S3**). Mean stiffness was 13 pN/nm (± 5 pN/nm, S.D.), which makes SARS-CoV-2 the most compliant virus investigated so far (Cieplak and Robbins, 2013; Mateu, 2012). Virion stiffness is somewhat lower than that measured for the influenza virus lipid envelope (Li et al., 2011), suggesting that the elasticity of SARS-CoV-2 is dominated by its envelope, and the RNP contributes little to the overall viral mechanics. The elastic regime was followed by a yield point marking the deviation from the linear force response and the onset of force-induced structural transitions which continued to take place until total compression. Unlike in other viruses (de Pablo, 2018; de Pablo and Schaap, 2019), force did not drop to near zero values following mechanical yield, indicating that virion collapse or breakage was not evoked in spite of the drastic mechanical perturbation. Conceivably, the mechanical yield is made possible by the force-induced rearrangement of the basketlike structures of the RNPs (Yao et al., 2020). Subsequent to indentation we retracted the cantilever. Remarkably, the virion generated forces up to several hundred pN during retraction, suggesting that structural recovery was taking place, possibly driven by the structural restitution of the initial RNP arrangement. The process continued until the initial virion height was reestablished. The mean force-spectroscopical height was 94 nm (± 10 nm, S.D.), which is comparable to that obtained from topographical data. The differences between the indentation and retraction force traces revealed a force hysteresis indicating that part of the mechanical energy invested in distorting the virion was dissipated as heat. We were able to continue the indentation-retraction cycles up to 100 times, but the virions never broke or collapsed (**Figs.3.e-f**). Rather, both the indentation and retraction force traces relaxed after about a dozen mechanical cycles, resulting in a minimized force hysteresis. The force response may potentially be explained by two other mechanisms alternative to virion compression. The first one is the force-induced virion rolling on the substrate. We exclude this possibility, because, due to the presence of anti-spike-protein antibodies on the surrounding surface, the process is expected to be completely irreversible. The second is the sideways slippage of the cantilever tip, off of the virion surface. We exclude this possibility based on a calibration of the cantilever’s lateral torsion (**Fig. S5**) and because this process is expected to be completely reversible. We can only speculate about the mechanisms behind the persistent structural self-healing of SARS-CoV-2. Conceivably, the process involves the dynamic interaction between the RNA, protein and lipid components. Notably, in some (~1 %) of the retraction traces we observed sawtooth-shaped force responses, the peak forces of which fall between ~210-330 pN (**Fig.3.f**). The most plausible explanation is that the force peaks correspond to the mechanically-driven unfolding of S protein domains (Moreira et al., 2020). Altogether, the SARS-CoV-2 virion is a mechanically stable, remarkably compliant and surprisingly resilient nanoparticle.

**Fig. 3.**
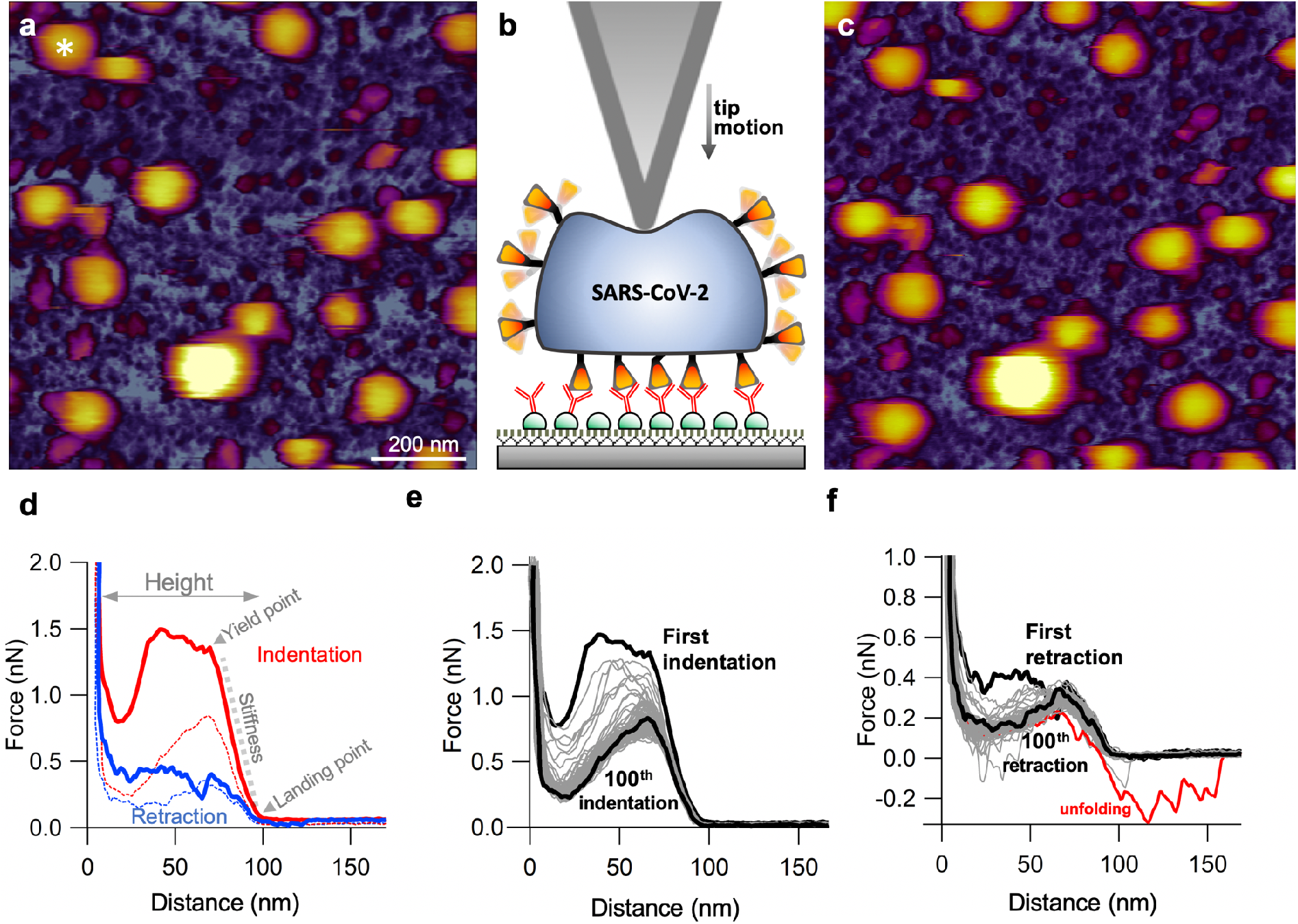
Single-particle force spectroscopy of SARS-CoV-2 virions. **a.** AFM image of an overview (1 x 1 μm) sample area prior to nanomechanical manipulation. Asterisk shows the the virion selected for nanomechanical manipulation. **b.** Schematics of the force-spectroscopy experiment. The virion is indented with the AFM tip until a pre-set force (typically 2-3 nN) is reached. **c.** AFM image of the same overview sample area following nanomechanical manipulation. We could not detect any topographical sign of permanent structural change. **d.** Example of a force *versus* distance curve obtained during a single indentation-retraction cycle. From the slope of the indentation curve (gray dotted line) and the distance between the landing point and substrate limit of the trace we obtained the stiffness and the force-spectroscopic height of the virions, respectively. Red and blue dotted lines indicate indentation and retraction data, respectively, obtained in the 100^th^ nanomechanical cycle. **e.** Force *versus* distance curves obtained during repeated indentation of a single SARS-CoV-2 virion. **f.** The matching force *versus* distance curves obtained during retraction. In some traces force sawteeth (red trace) corresponding to the unfolding of domains in a surface protein, plausibly within the S trimer.

### Thermal sensitivity of SARS-CoV-2

To assess the thermal stability of SARS-CoV-2, we explored the topographical changes of virions exposed to high-temperature treatment (**Fig.4**). The sample was exposed to temperatures of 60 (**Fig.4.a**), 80 (**Fig.4.b**) and 90 °C (**Fig.4.c**) for ten minutes then cooled back to 20 °C for AFM imaging. Remarkably, virions remained on the substrate surface, and their global appearance was only slightly altered. Virion topography became somewhat faceted, but the particles retained their blurred, smooth surface. To test whether spikes were still present following thermal exposure, we fixed the sample with 5% glutaraldehyde (**Fig.4.d**). Although the rugged topography, seen in chemically fixed SARS-CoV-2 virions (**Fig.1.c**), was partially restored, the spikes were much fewer, less distinct, and their trigonal shape (**Fig.1.e**) could not be resolved (**Fig.4.e**), suggesting they became thermally denatured. Furthermore, the smooth areas interspersed between rugged regions indicate that thermal treatment resulted in a progressive dissociation of the S trimers from the virion surface. Thus, the SARS-CoV-2 virion displays an unexpected global thermal stability, which is likely related to their aerosol and surface stabilities (van Doremalen et al., 2020). However, the conformational response of the spike proteins observed here eventually leads to the heat-induced inactivation of SARS-CoV-2.

**Fig. 4.**
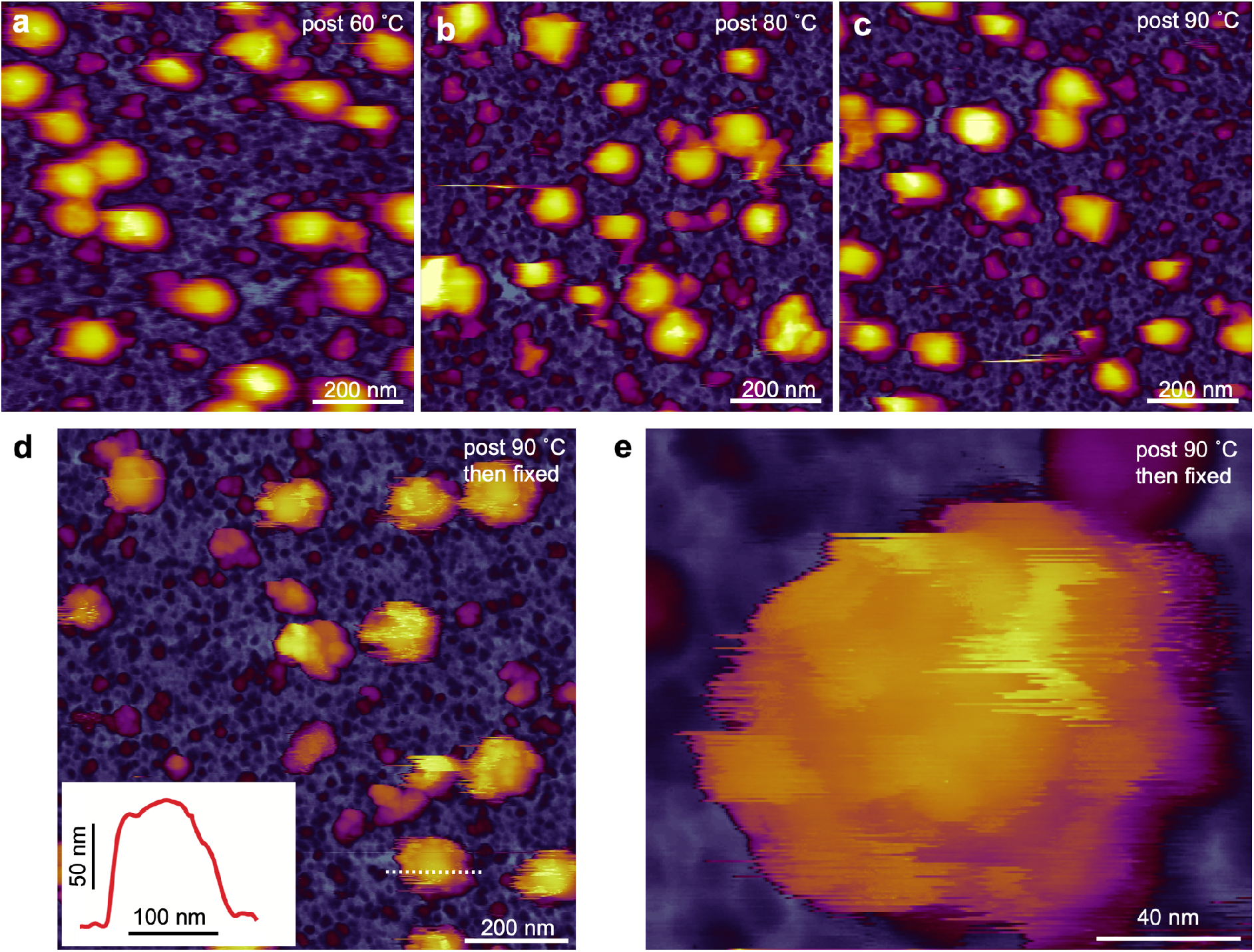
Effect of temperature change on the topographical structure of SARS-CoV-2. The sample was heated for ten minutes at 60 (**a**), 80 (**b**) and 90 °C (**c**), then cooled back to 20 °C prior to AFM imaging. The virions persist on the substrate surface with their global structure nearly intact, but the topography becomes progressively more rugged, pointing at the gradual disappearance of the dynamic surface smoothing hence reduction of spike dynamics. **d.** AFM image of an overview (1 x 1 μm) sample area following thermal treatment (90 °C for ten minutes) and glutaraldehyde (5%) fixation. **Inset**, topographical profile plot measured along the center of one of the virions (dotted line). The rugged surface topography is partially restored, but large areas on the virions are devoid of spikes. **e.** High-resolution AFM image of a heat-exposed (90 °C for ten minutes) and fixed (5% glutaraldehyde) SARS-CoV-2 virion. Shallow surface protrusions are present.

## CONCLUSIONS

The atomic force microscopic imaging and nanomechanical measurements revealed that the SARS-CoV-2 virion is highly dynamic, compliant and resilient, and it displays remarkable mechanical and global thermal stabilities. While the dynamics of the surface spikes may play an important role in the unusually high infectivity of the virus, its mechanical and self-healing properties may also ensure adaptation to a wide range of environmental circumstances. Considering its capability of exploring viruses under native conditions, the single-particle approaches employed here may be important in uncovering not only the mechanistic details behind viral infection but the viral response to potential therapies as well.

## ACKNOWLEDGEMENTS

We gratefully acknowledge the help of Mónika Drabbant and Hedvig Tordai with experimental preparations and measurements. We thank Levente Herényi for theoretical discussions and reading the manuscript. This work was funded by grants from the Hungarian National Research, Development and Innovation Office (K124966 to MK and FK128956 to ZM; National Heart Program NVKP-16-1-2016-0017; Thematic Excellence Programme; National Bionics Programme ED_17-1-2017-0009). The research was financed by the Higher Education Institutional Excellence Programme of the Ministry for Innovation and Technology in Hungary, within the framework of the Therapeutic Development thematic programme of the Semmelweis University.

## AUTHOR CONTRIBUTIONS

B.K., Z.K., B.P. and M.S.Z.K. conceived experiments; Z.K. and B.P. purified the SARS-CoV-2 samples; B.K. and M.S.Z.K. performed AFM imaging and force spectroscopy measurements and analyzed data; B.K. and M.S.Z.K. wrote paper; M.S.Z.K. secured financial support for the project. All authors critically read and revised the manuscript.

## AUTHOR DECLARATIONS

The authors declare no conflict of interest.

## MATERIALS AND METHODS

### Sample preparation

SARS-CoV-2 was isolated from the oropharyngeal swab of a laboratory-confirmed COVID-19 patient in Hungary and passed two times in VeroE6 cell line (European Collection of Authenticated Cell Culture, Salisbury, U.K.) in Dulbecco’s Modified Eagle’s Medium (Lonza, Basel, Switzerland) supplemented with 5% fetal bovine serum (EuroClone, Pero, Italy) and Cell Culture Guard (PanReac AppliChem, Darmstadt, Germany). To remove the disturbing effects of fetal bovine serum albumin, an additional passage was carried out in VP-SFM serum-free, ultra-low protein medium (Gibco, ThermoFisher Scientific) supplemented with L-glutamine (Sigma-Aldrich, Merck, Darmstadt, Germany). Four days after inoculation, when full cytopathic effects were observed, the virus-containing medium was collected and centrifuged (3000 x g) to remove debris. To concentrate the virus, the supernatant was ultracentrifuged (70,000 x g, 1.5 hours, 4°C) in 13.5-ml lockable plastic tubes using a Sorvall MTX-150 ultracentrifuge. The supernatant was removed and the pellet was resuspended in 100 μl VP-SFM. All sample preparation steps were performed in biosafety level-3 (BSL-3) conditions.

### Preparation of affinity-enhanced substrate surface

100 μl of 0.1% w/v poly-L-lysine (PLL) (Merck, Darmstadt, Germany) solution was pipetted onto a freshly cleaved mica surface (Ted Pella, Redding, CA) and incubated for 20 minutes, following which the surface was rinsed with distilled water and dried in N2 stream. 100 μl of 25% w/v grade I glutaraldehyde (GA) (Merck, Darmstadt, Germany) was then pipetted on the surface and incubated for 30 min, followed by further rinsing with distilled water and drying in N2 stream. Subsequently, 100 μl of 10 μg/ml recombinant protein G (Merck, Darmstadt, Germany) was added and incubated for 30 minutes, then the surface was washed five times each with 100 μl phosphate-buffered saline (PBS). 100 μl of 10 μg/ml SARS-CoV-2 Spike Glycoprotein Antibody (#abx376478, Abbexa Ltd, Cambridge, UK) was then added, and the surface was incubated for 1 hour. All surface functionalization steps were performed at room temperature. Unbound antibodies were removed by repeated rinsing with PBS. The anti-spike glycoprotein antibody-coated surfaces were stored under PBS for up to 5 days at 4 °C.

### Handling virion sample in the AFM

An aliquot (20 μl) of purified SARS-CoV-2 sample was pipetted onto the anti-spike antibody-coated substrate surface and incubated at 37 °C for 30 minutes. To increase virion surface density, another aliquot was added and incubated. The process was repeated twice. Subsequently, the surface was rinsed gently with PBS to remove unbound virions. All the sample-loading and washing steps were carried out in a laminar-flow hood in BSL-3 conditions (at the National Biosafety Laboratory, National Public Health Centre, Hungary). For AFM imaging of chemically fixed SARS-CoV-2, 100 μl 5% GA solution (in PBS) was added, and the sample was incubated for >1 hour, ensuring both fixation and virus inactivation. Then, the sample was carried to the AFM laboratory (Department of Biophysics and Radiation Biology, Semmelweis University) for loading in the environmental scanner unit of the Cypher ES AFM instrument. For imaging native virions, the fixation step was omitted and the sample was loaded directly in the Cypher ES Scanner. To ensure compliance with safety measures, a closed cantilever holder and gas-tight sample chamber was used, and the sample was loaded into the scanner in a laminar-flow hood in BSL-3 conditions. Then the scanner was carried to the AFM laboratory and inserted into the AFM instrument for imaging. To further comply with safety measures, following AFM imaging the native virus samples were discarded in either of two ways. In the first, the AFM scanner was taken to the BSL-3 laboratory for removal and chemical desctruction of the sample. In the second, the sample was heated to 90 °C for >10 minutes with the temperature-controller unit of the Cypher ES scanner. Then, the sample was removed for immediate chemical destruction (5% NaClO).

### AFM imaging and force spectroscopy

Atomic force microscopy imaging was carried out with an Asylum Research Cypher ES instrument (Oxford Instruments, Santa Barbara, CA)(Kellermayer et al., 2018; Voros et al., 2017). Resonant-mode (AC- or non-contact mode) scanning was performed under liquid in PBS with BL-AC40TS (Olympus Corporation, Japan) cantilevers. The cantilever was oscillated near its resonance frequency (tyically around 20 kHz) by using photothermal excitation at a typical free amplitude of 0.5 V. Imaging was carried out at a typical setpoint of 350 mV and with scanning speeds <1 μm/s to prevent the mechanical dislodging of virions from the substrate surface. Alternatively, we used fast force mapping (FFM, or “jumping-mode” AFM(Moreno-Herrero et al., 2004)) to collect image data. In FFM the cantilever was driven sinusoidally with a typical frequency of 300 Hz and a setpoint force of 100 pN to obtain a force curve for each pixel, from which the topographical image was reconstructed. The two (AC-mode and FFM) imaging modes gave similar results. Thermal exposure of the virions was achieved by heating the sample stage to pre-set temperatures(Voros et al., 2018). Force spectroscopic measurements were carried out on virions selected on the AFM images. The cantilever was lowered, in contact mode, onto the vertex of the virion with 0.5 μm/s velocity until the setpoint force was reached (typically 2-3 nN). Cantilever spring constants were determined by the thermal method(Hutter and Bechhoefer, 1993) prior to imaging. Spring constants were between 90-120 pN/nm.

### Data analysis

Image postprocessing and data analysis were performed by using the AFM driving software AR16, IgorPro 6.37 (Wavemetrics, Lake Oswego, OR). Particle analysis was carried out in subsequent analytical steps. First, the particles were demarcated by masking at the full width at half maximal height. Second, the mask was eroded, then dilated to gently smooth particle edges, while also getting rid of small scanning artefacts. We ignored particles with extreme deformities, ones too close to each other or with an area smaller than 3000 nm^2^. Height was calculated at the center of the particles. Volume was calculated as the sum of the heights of individual pixels of the particle multiplied by the spatial (x, y) scaling values to correct for image pixel resolution. Particle diameter was calculated as the diameter of a circle with the same area as the particle itself.

