## Supplementary material for "Topography, spike dynamics and nanomechanics of individual native SARS-CoV-2 virions"

[#]These authors contributed equally

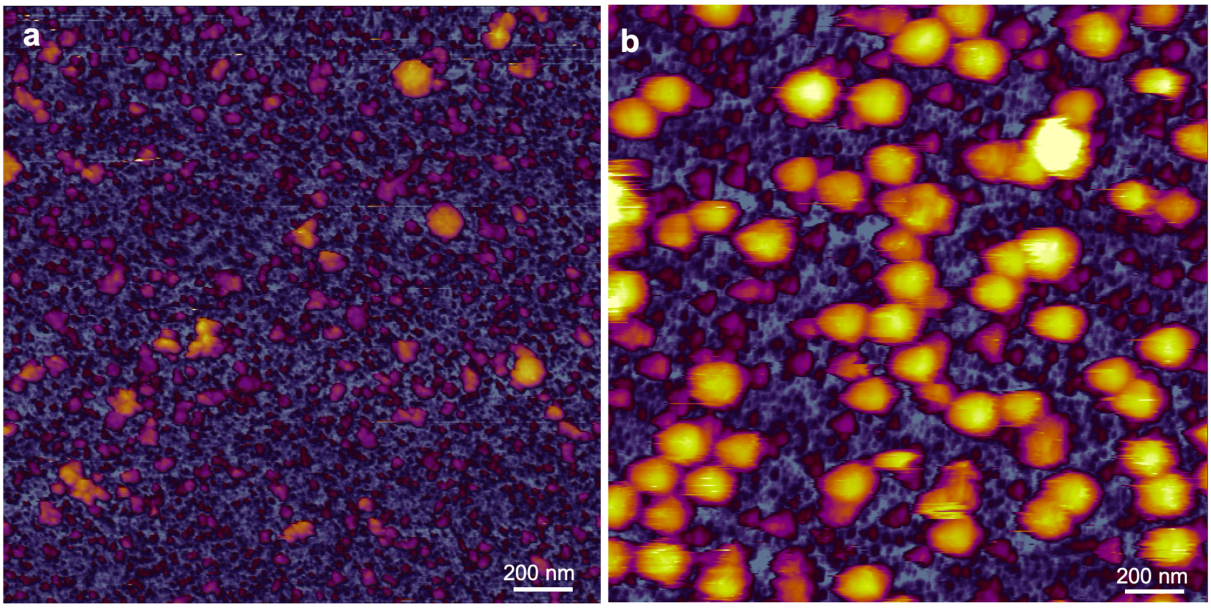


**Fig. S1.** Effect of using monoclonal anti-spike antibody for affinity-enhanced surface binding of SARS-Cov-2 virions. **a.** AFM image of an overview (2 x 2 μm) sample area where protein G and the anti-spike protein antibody were omitted, and the virus sample was directly loaded onto a mica surface coated with poly-L-lysine and glutaraldehyde. **b.** AFM image of an overview (2 x 2 μm) sample area where the viruses were captured by the anti-spike protein antibody on the substrate surface. Binding was amplified by nearly two orders of magnitude.

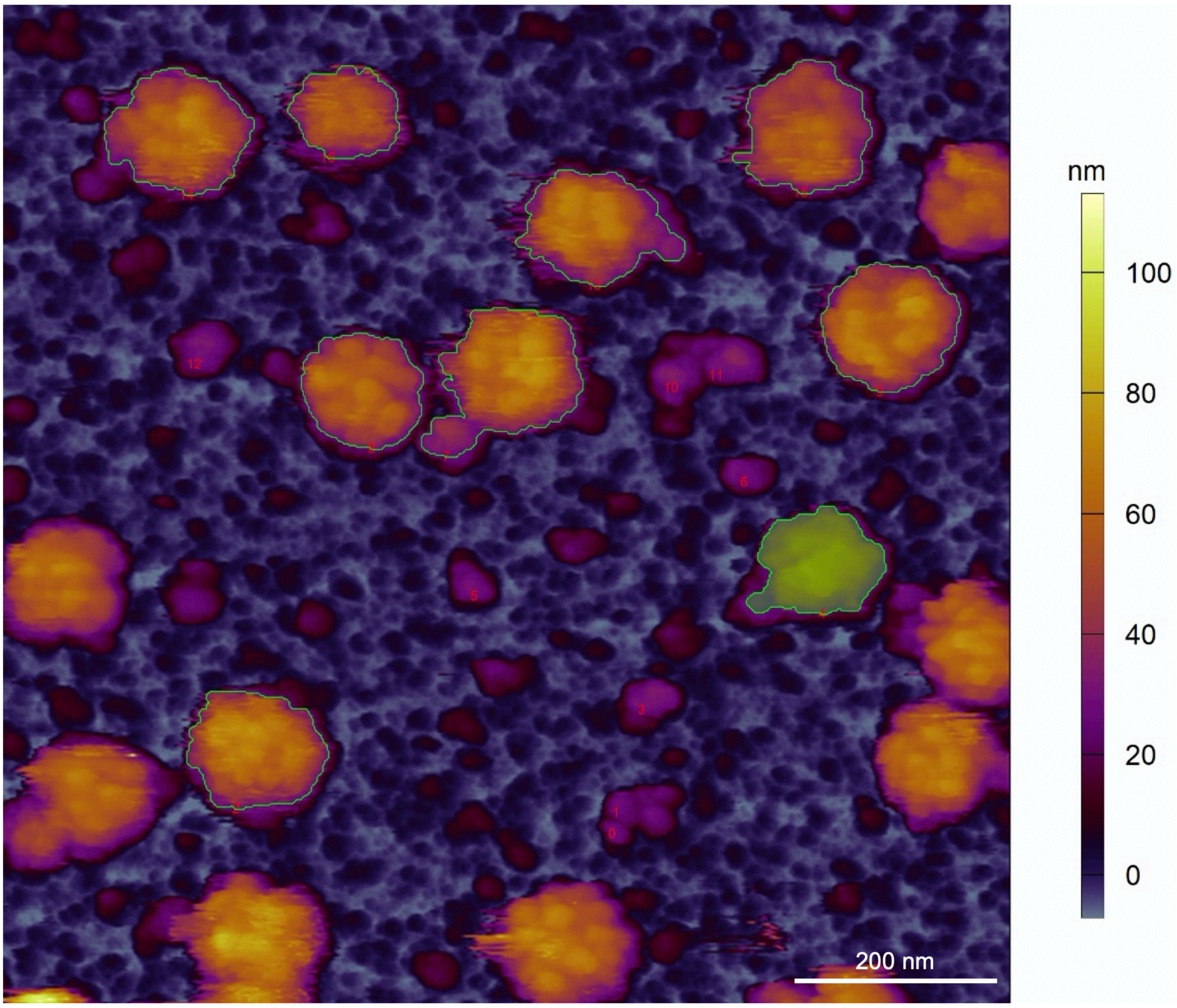


**Fig. S2.** Particle analysis carried out in AFM images of SARS-CoV-2 virions. Green line surrounding particles shows the contour at the average half peak height, which defines the border of the analyzed particles. One such particle fit for analysis is filled with green colour.

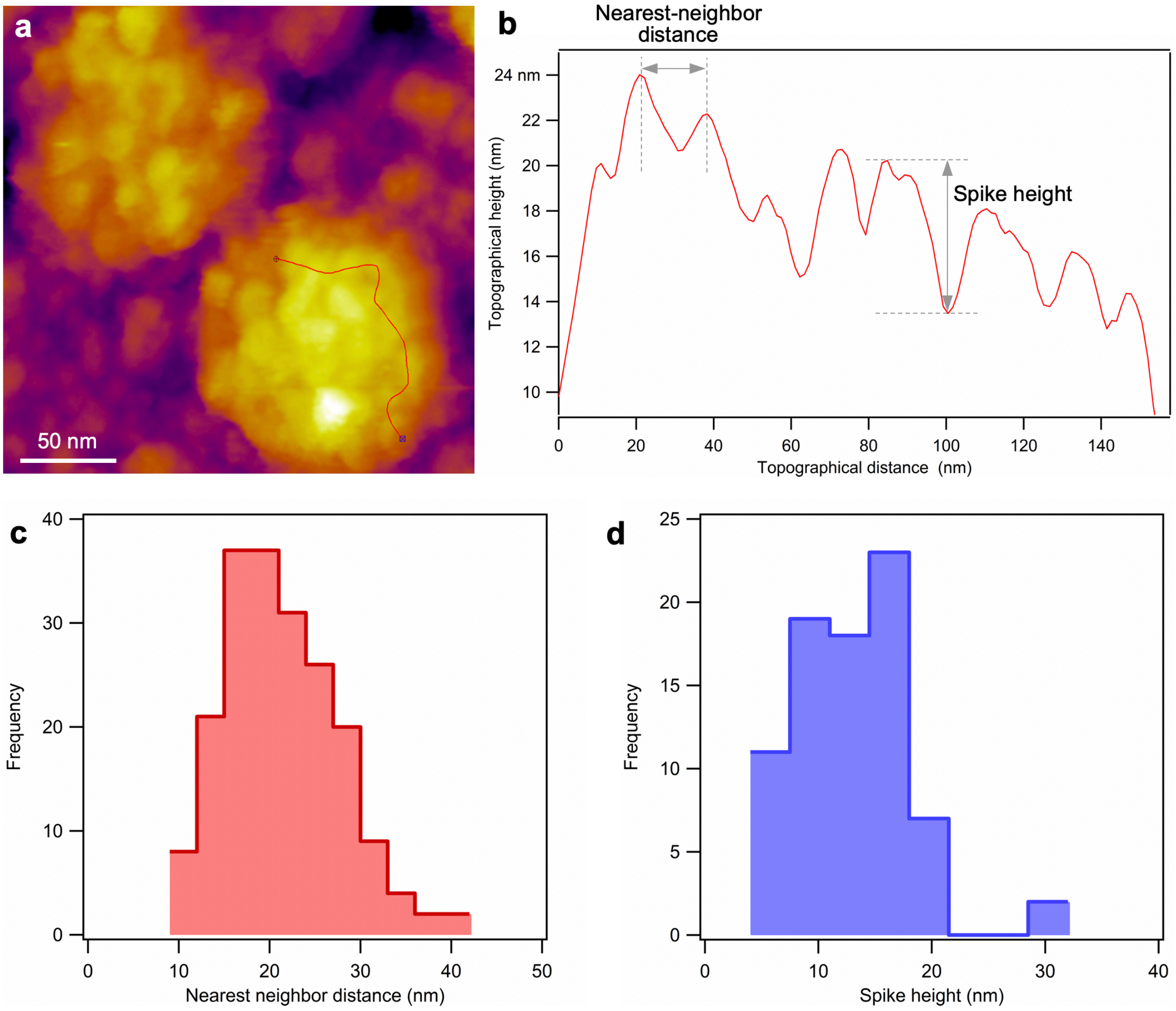


**Fig. S3.** Measurement of the nearest-neighbor distance between and the height of S trimers. **a.** AFM image of two SARS-CoV-2 virions fixed with glutaraldehyde (5%). Red line indicates the path, drawn approximately in coronal plane, along which the topographical height was measured. **b.** Topographical profile plot of the virion along the line indicated in **a**. Nearest-neighbor distance was obtained by measuring the distance between subsequent peaks. Spike height was obtained by measuring the distance, along the height axis, between a peak and the following valley. Height was measured only for spikes with a large enough valley in their vicinity so as to increase the probability of the AFM tip reaching the envelope surface. **c.** Distribution of the nearest neighbor distance between spikes (n=197). **d.** Distribution of the spike height (n=80).

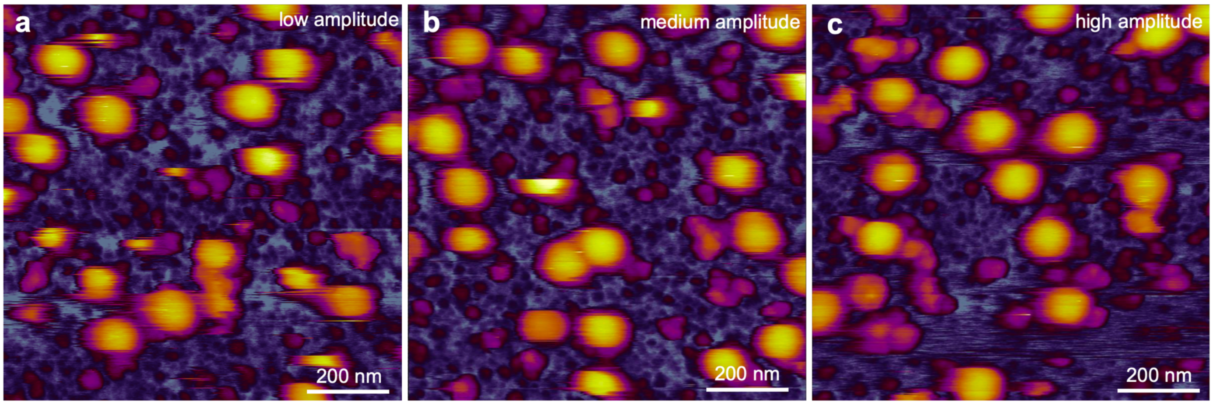


**Fig. S4.** Effect of cantilever oscillation amplitude on imaging native, unfixed SARS-CoV-2 virions. Setpoint per free-amplitude values were 120 mV/155 mV (**a**), 210 mV/280 mV (**b**) and 385 mV/500 mV (**c**).

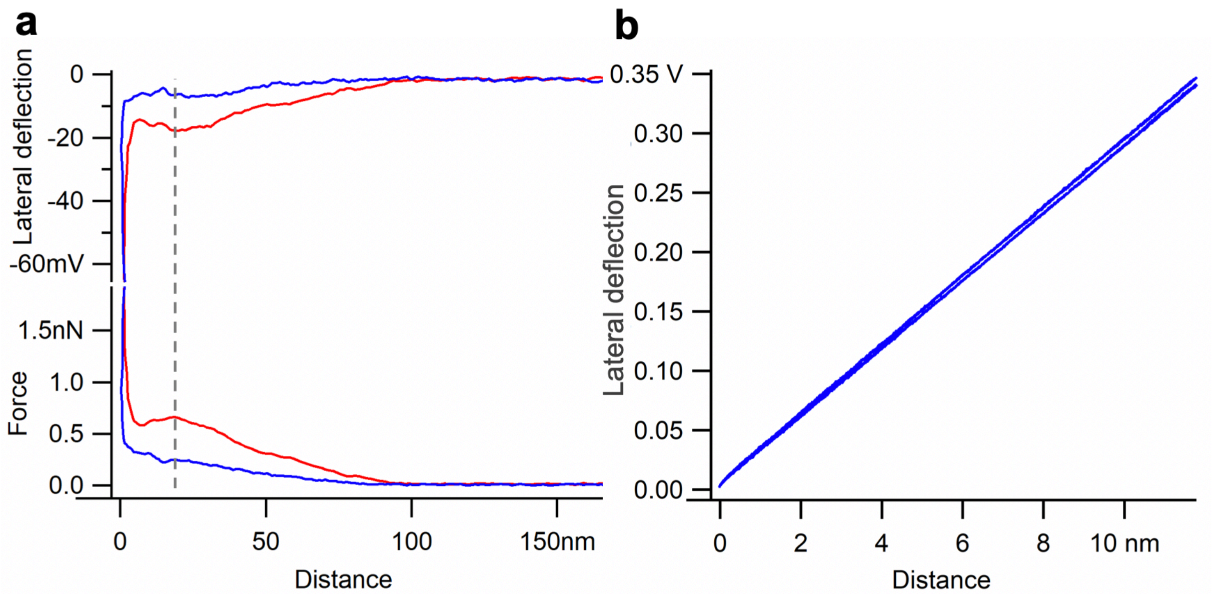


**Fig. S5.** Testing the contribution of cantilever torsion to the force spectrum. **a.** Force (in nN) and lateral displacement signal (in mV) as a function of distance obtained during an indentation-retraction experiment on a single SARS-CoV-2 virion. **b.** Calibration of lateral displacement. Lateral displacement (in volts) as a function of distance is shown. The trace was obtained in an experiment in which the cantilever was pressed, with a force of 2 nN, into a mica surface so as to immobilize the tip, then the cantilever was moved laterally (along a direction perpendicular to its long axis) with an amplitude of 10 nm. The nearly overlapping blue traces correspond to data acquired during back-and-forth motion. Such an experiment resulted in the torsion of the cantilever and provided means to measure the signal (volts) per unit torsional displacement (nm). From the slope of the line (29 mV/nm) we obtained the lateral distance calibration. The calibration indicated that the 19 mV lateral displacement signal measured at the 0.7 nN indentation force (gray dotted line in **a**) corresponds to a lateral distance of only 6.5 Å. Because this small distance is negligible in comparison to the virion diameter (<1 %), we exclude the possibility that during indentation the cantilever slips sideways off the virus particle.

### SUPPLEMENTARY TABLES

**Table S1.** Particle analysis results. Particle height was measured in the center of particles. We used the mean height of the native virion ($d_v$) to calculate its mean surface area ($A_v$) as $d_v^2\pi$ = 21642 $nm^2$.

| | Height (nm) | Volume ($nm^3$) | Diameter (nm) | n |
|---|---|---|---|---|
| Fixed | 62 ± 8 | 574 000 ± 212 000 | 120 ± 16 | 51 |
| Native | 83 ± 7 | 490 000 ± 107 000 | 99 ± 11 nm | 47 |
| Heated (90 ˚C) | 82 ± 10 | 600 000 ± 152 000 | 108 ± 12 | 37 |

**Table S2.** Spike analysis results. We assume that each spike occupies a circular area on the virion surface the diameter ($d_s$) of which is the mean nearest-neighbor distance. Hence, the mean area occupied by a spike is $(d_s/2)^2\pi$ = 356 $nm^2$. The ratio of the virion and spike-occupancy surfaces provides the average number of spikes on a virion: ~61.

| | Mean ± S.D. (nm) | n |
|---|---|---|
| Nearest-neighbor distance | 21 ± 6 | 197 |
| Spike height | 13 ± 5 | 80 |

**Table S3.** Structural and nanomechanical data extracted from force spectroscopy results.

| | Height (nm) | Stiffness (pN/nm) | n |
|---|---|---|---|
| Native virion | 94 ± 10 | 13 ± 5 | 40 |
